# Pore-forming protein βγ-CAT drives extracellular nutrient scavenging under cell starvation

**DOI:** 10.1101/2022.07.20.500773

**Authors:** Ling-Zhen Liu, Long Liu, Zhi-Hong Shi, Xian-Ling Bian, Qi-Quan Wang, Yang Xiang, Yun Zhang

## Abstract

Nutrient acquisition is essential for cells. βγ-CAT is a pore-forming protein (PFP) and trefoil factor complex assembled under tight regulation identified in toad *Bombina maxima*. Here, we reported that *B. maxima* cells secreted βγ-CAT under glucose and glutamine deficiency to scavenge extracellular proteins for their nutrient supply and survival. AMP-activated kinase signaling positively regulated the expression and secretion of βγ-CAT. The PFP complex promoted albumin and ovalbumin uptake through endolysosomal pathways. Elevated intracellular amino acids, enhanced ATP production, and eventually prolonged cell survival were observed in the presence of βγ-CAT and extracellular albumin or ovalbumin. Liposome assays indicated that high concentration of ATP (around 1–5 mM) negatively regulated the opening of βγ-CAT channels. Collectively, these results uncovered that βγ-CAT is an essential element in cell nutrient scavenging under cell starvation by driving vesicular uptake of extracellular proteins, providing a new paradigm for PFPs in cell nutrient acquisition and metabolic flexibility.

## Introduction

Plasma membrane transporters and receptor-mediated endocytosis of nutrient carriers are the two main canonical ways by which cells import nutrients for their life cycle. The former operates in uptake of small nutrient, such as glucose and amino acids, and the latter is the main pathway through which cells obtain insoluble nutrients, such as cholesterol and iron (1, 2). Additionally, cells ingest extracellular macromolecules through endolysosomal systems via macropinocytosis, an evolutionarily conserved form of endocytosis mediating non-selective import of fluid and solutes contained therein (3-5). Cells import macromolecules such as proteins and degrade them in lysosomes to support their metabolism and growth, such as in cancer cell metabolism (5, 6). The regulation and coordination of distinct nutrient acquisition strategies are incompletely understood (2, 7, 8). The possible existence and identity of extracellular elements dispatched by cells for nutrient sampling and scavenging under starvation require exploration.

Numerous pore-forming proteins (PFPs) with a membrane insertion domain similar to bacterial toxin aerolysin, namely aerolysin family PFPs (af-PFPs, previously referred as aerolysin-like proteins, ALPs), have been found in plants and animals (9-11). BmALP1 is an af-PFP from toad *B. maxima*. It forms membrane pores (channels) with a functional diameter 1.5–2.0 nm (12-14). This PFP is regulated by environmental cues and interacts with a trefoil factor (BmTFF3) to form an PFP complex βγ-CAT, in which BmTFF3 acts as a chaperon and a regulatory unit of BmALP1 (14-17). BmALP3, a paralog of BmALP1, lacks a membrane pore-forming capacity, but it oxidizes BmALP1 to its water-soluble polymer, leading to the dissociation of βγ-CAT complex and loss of biological activity (17).

This PFP complex targets cell surface acidic glycosphingolipids in lipid rafts via a double-receptor binding model (16) to act along cell endocytic and exocytic pathways with channel formation on endolysosomes, which have been shown to play roles in immune defense and tissue repair (18-22). Thus, the PFP and its regulatory network define an unknown secretory endolysosomal channel (SELC) pathway, representing a novel PFP-driven cell vesicular delivering system (13, 23). The newly defined βγ-CAT pathway is able to mediate cell import and export of extracellular materials and/or plasma membrane components through endolysosomal systems, making the PFP complex a versatile multiple functional protein machine depending on the cell context and surroundings. βγ-CAT drives macropinocytosis to facilitate water maintaining in toad osmoregulatory organs in response to osmotic stress (13). Toad blood βγ-CAT is an immediate responsive element under animal fasting, which has been proposed to mediate transcellular transport of albumin-bound fatty acids for nutrient supply of tissue parenchymal cells (24). However, the possible involvement of the PFP complex in cell responses to starvation and its specific roles in starved cells remain unclear.

Here, we used the starvation model of toad *B. maxima* cells to further investigate the role of SELC protein βγ-CAT in cell nutrient acquisition and metabolic flexibility under cell nutrient deficiency. Interestingly, toad liver and gastrointestinal cells secreted βγ-CAT to scavenge extracellular protein nutrients, such as albumin and ovalbumin, through endolysosomal pathways in the absence of glucose and glutamine, which elevated intracellular amino acids and ATP levels and supported cell survival under starvation. The expression and secretion of βγ-CAT were largely attenuated by inhibition of AMP-activated kinase (AMPK) signaling. Furthermore, in a liposome model, ATP, but not AMP, inhibited βγ-CAT channel opening. These results revealed the essential role of a secretory PFP in extracellular nutrient scavenging by cells through endolysosomal pathways under cell starvation.

## Results

### Toad cells secrete βγ-CAT under nutrient deficiency

Previous studies have shown that secreted βγ-CAT promotes cell import through endolysosomal pathways probably via inducing pinocytosis/macropinocytosis in distinct cell context (13, 21, 24). This raises the possibility that the PFP protein complex participates in macromolecule intake for cellular nutrient supply and metabolic flexibility under variations in nutrient availability (13, 23, 24). To test this hypothesis, we used a starvation model of toad cells cultured under three nutrient conditions including glucose/glutamine-containing medium (Glc^+^/Gln^+^), glucose/glutamine-depleted medium (Glc^-^/Gln^-^), and glucose-containing but glutamine-depleted medium (Glc^+^/Gln^-^). In the isolated toad liver cell population, there were 62.4% hepatocytes (Fig. S1A) as assessed by a specific antibody against hepatocyte marker cytokeratin 18 (CK18) (25, 26). Under Glc^-^/Gln^-^ conditions, the expression of βγ-CAT α-subunit BmALP1 was attenuated at 1 hour, but substantially upregulated at 3 hours and 5 hours as analyzed by qRT-PCR (Fig. 1A) and western blotting (Fig. 1B). Meanwhile, the expression of βγ-CAT β-subunit BmTFF3 was also upregulated at 3 hours (Fig. 1A). Because the PFP complex βγ-CAT is a secreted protein, we next investigated the change in the βγ-CAT protein level in toad liver cell supernatants by blotting for its α-subunit BmALP1. Under Glc^-^/Gln^-^ conditions, largely augmented secretion of βγ-CAT α-subunit BmALP1 was readily detected by western blotting (Fig. 1C). Biologically active βγ-CAT in culture supernatants was analyzed by its hemolytic activity on human erythrocytes, a sensitive method to determine the existence of βγ-CAT (13, 17). First, we verified that the various culture media did not affect the hemolytic activity of βγ-CAT (Fig. S1B). Intriguingly, the hemolytic activity in culture supernatants of toad liver cells was largely increased under Glc^-^/Gln^-^ conditions compared with that of Glc^+^/Gln^+^ conditions (Fig. 1D), which was abolished by anti-βγ-CAT antibodies (Fig. S1C). These results indicated secreted βγ-CAT in culture supernatants of liver cells under starvation.

**Figure 1.**
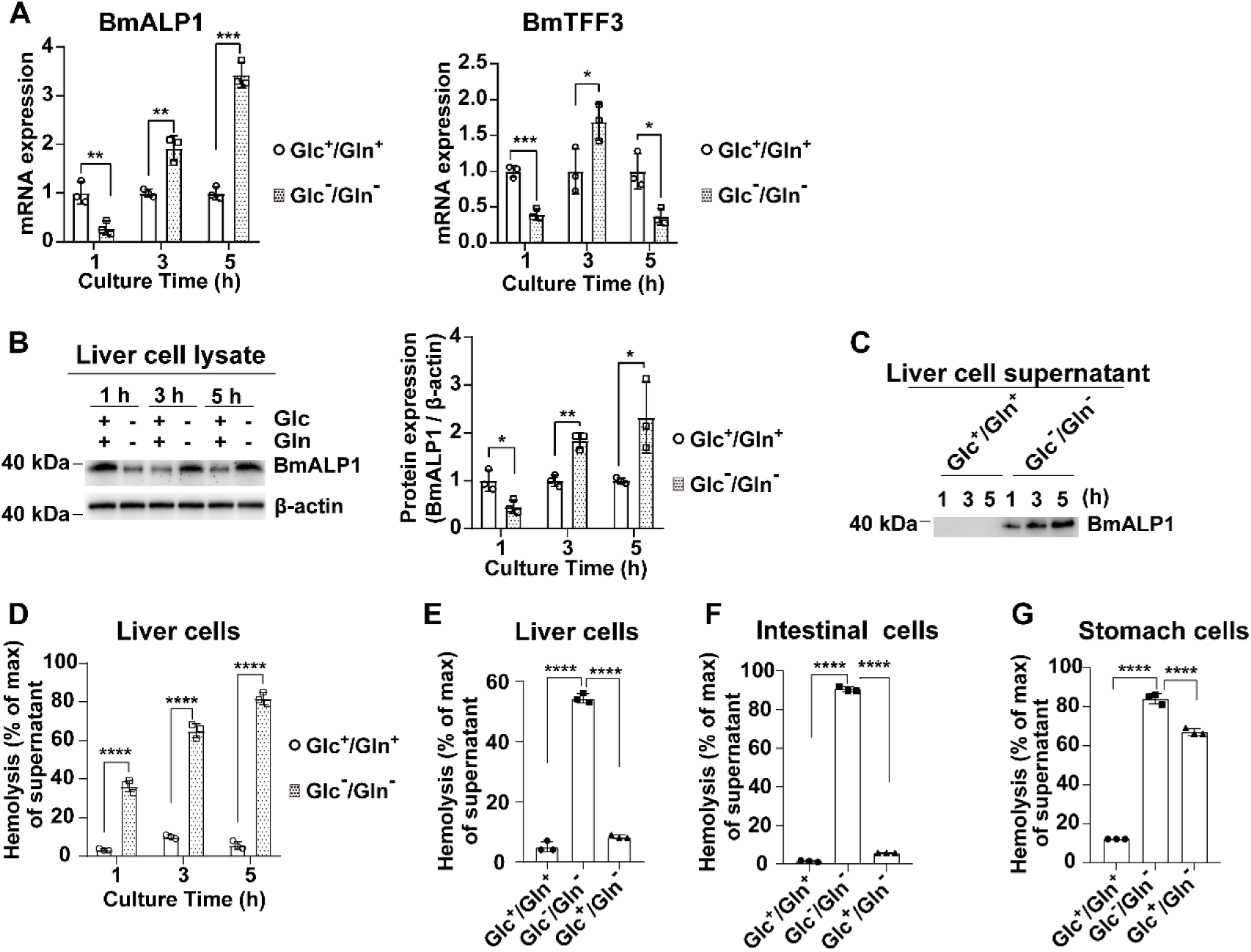
Expression and secretion of βγ-CAT are substantially enhanced in cell response to nutrient deficiency. (A–D) Primary isolated toad liver cells were cultured in Glc^+^/Gln^+^ medium or starved in Glc^-^/Gln^-^ medium for 1–5 hours. (A) BmALP1 (*Left*) and BmTFF3 (*Right*) mRNA levels were determined by qRT-PCR. (B) The protein level of βγ-CAT α-subunit BmALP1 in cell lysates was determined by western blotting (*Left*) and the bands were semi-quantified by ImageJ (*Right*). (C, D) Secretion of βγ-CAT in liver cell supernatants was determined by western blotting (C) and a hemolytic activity assay (D). (E–G) Toad cells were cultured in Glc^+^/Gln^+^, Glc^-^/Gln^-^, or Glc^+^/Gln^-^ medium for 1 hour (intestinal and stomach cells) or 3 hours (liver cells). Hemolytic activity was detected in liver cell supernatants (E), intestine cell supernatants (F), and stomach cell supernatants (G). Data (A, B and D–G) are represented as the mean ± SD of triplicate samples. \**P*<0.05, \*\**P*<0.01, \*\*\**P*<0.001, and \*\*\*\**P*<0.0001 by unpaired *t-*test (A, B, D) or one-way ANOVA (E– G). All data are representative of at least two independent experiments.

As the primary energy source, glucose is rapidly used by cells for energy supply (27). Next, we investigated secretion of βγ-CAT under Glc^+^/Gln^-^ conditions. The hemolytic activity assay of culture supernatants showed that the secretion of βγ-CAT was substantially decreased in the liver cell supernatants under Glc^+^/Gln^-^ conditions as compared with that of Glc^-^/Gln^-^ conditions (Fig. 1E), indicating that the presence of glucose largely decreased βγ-CAT secretion in toad liver cells. A similar result was observed in toad intestinal cells (Fig. 1F). However, the secretion of βγ-CAT was only partially attenuated under Glc^+^/Gln^-^ conditions in toad stomach cells under our assay conditions (Fig. 1G). These results showed that toad cells from the alimentary system secrete βγ-CAT to counteract nutrient deficiency, and glutamine depletion alone could also result in secretion of the PFP complex.

Taken together, these results revealed that the expression and secretion of βγ-CAT are substantially upregulated under cell nutrient deficiency, and the PFP protein complex is an immediate responsive protein machine to cell starvation. This phenomenon is well in accordance with our previous observation that the PFP complex in toad blood promptly responds to toad fasting (24).

### AMPK signaling positively regulates the expression and secretion of βγ-CAT under cell starvation

AMPK signaling is activated by a lack of energy or nutrients and switches on alternative catabolic pathways that generate ATP while switching off anabolic pathways and other processes that consume ATP (28, 29). Because toad cells secrete βγ-CAT under nutrient deficiency (Fig. 1), the expression and/or secretion of the PFP complex might be controlled by AMPK signaling.

Sequence alignment analysis of toad *B. maxima* AMPK α-subunits and acetyl coenzyme A carboxylase 1 (ACC1) on the basis of toad skin transcriptome (30) verified that the activation loop of AMPKs and phosphorylation sites (pAMPK^T172^ and pACC1^S79/80^, respectively) were evolutionarily conserved from toad *B. maxima* to human (Fig. S2A). Activation of AMPK signaling in toad liver cells was observed under starvation (Fig. S2B). Thus, we used two pharmacological AMPK signaling inhibitors, compound C and SBI-0206965 that act on the activation loop of AMPK α-subunit (31, 32), to examine the possible effects of AMPK signaling on βγ-CAT regulation under cell starvation. We first analyzed the cytotoxicity of these AMPK signaling inhibitors in toad liver cells. The results showed that dosages up to 10 μM compound C and 20 μM SBI-0206965 did not affect the viability of toad liver cells (Fig. S2C and S2D). The presence of compound C (2.5–5 μM) or SBI-0206965 (5–10 μM) did inhibit the activation of AMPK signaling in toad liver cells as observed by reduced phosphorylation of the canonical AMPK substrate ACC1 under Glc^-^/Gln^-^ conditions (Fig. S2E).

After validating the inhibitory effects of compound C and SBI-0206965 on AMPK signaling in toad liver cells, we next analyzed whether the expression and secretion of βγ-CAT were regulated by AMPK signaling under cell starvation. The mRNA levels of βγ-CAT subunits BmALP1 and BmTFF3 were substantially decreased in the presence of compound C (Fig. 2A) or SBI-0206965 (Fig. 2E) under Glc^-^/Gln^-^ conditions. Moreover, the protein level of βγ-CAT, as indicated by detecting its α-subunit BmALP1, was largely decreased in toad liver cells after treatment with 5 μM compound C (Fig. 2B) or 10 μM SBI-0206965 (Fig. 2F) for 3 hours under Glc^-^/Gln^-^ conditions. Importantly, the treatment of toad liver cells with these two AMPK inhibitors substantially reduced βγ-CAT secretion into the culture supernatant of the toad cells. Western blotting showed greatly reduced secretion of BmALP1 (α-subunit of βγ-CAT) in culture supernatants of toad liver cells under Glc^-^/Gln^-^ conditions (Fig. 2C and 2G). The decreased hemolytic activity of the culture supernatants further confirmed that βγ-CAT secretion was attenuated by these AMPK signaling inhibitors (Fig. 2D and 2H). Therefore, these results revealed that AMPK activation controls the expression and secretion of βγ-CAT under glucose and glutamine starvation as the main carbon source.

**Figure 2.**
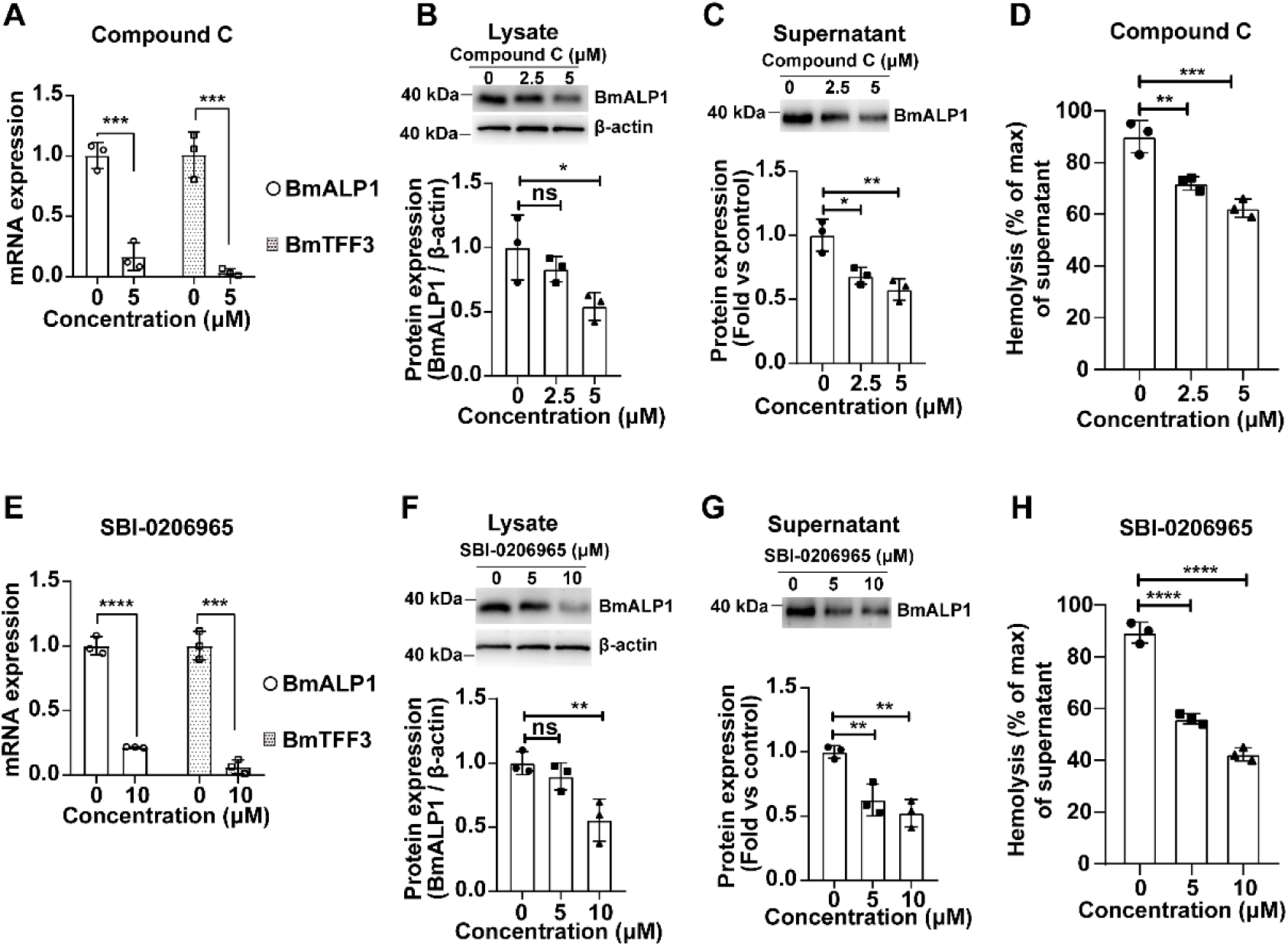
Expression and secretion of βγ-CAT are positively regulated by AMPK signaling. Isolated toad liver cells were treated with AMPK inhibitors compound C (A–D) or SBI-0206965 (E–H) in Glc^-^ /Gln^-^ medium for 3 hours. BmALP1 and BmTFF3 mRNA levels (A, E) were determined by qRT-PCR. The protein levels of βγ-CAT α-subunit BmALP1 in cell lysates (B, F) and supernatants (C, G) were detected by western blotting (*Top*) and the bands were semi-quantified by ImageJ (*Bottom*). The AMPK inhibitor was omitted as a control in the quantification shown in (C, G). The hemolytic activity (D, H) of culture supernatants was detected by hemolytic activity assay. Data are represented as the mean ± SD of triplicate samples. ns (*P*≥0.05), \**P*<0.05, \*\**P*<0.01, \*\*\**P*<0.001, and \*\*\*\**P*<0.0001 by the unpaired *t-* test (A, E) or one-way ANOVA (B–D, F–H). All data are representative of at least two independent experiments.

### βγ-CAT promotes extracellular protein import to facilitate intracellular amino acid supply and ATP production

Because toad cells dispatch the PFP complex βγ-CAT to the extracellular medium under starvation downstream AMPK signaling (Figs. 1 and 2), we next explored the possible cellular functions of βγ-CAT under cell nutrient deficiency. It has been proposed that βγ-CAT represents a novel PFP-driven cell vesicular delivery system (13, 23, 24), which has been shown to stimulate cell pinocytosis/macropinocytosis to promote cellular material import including extracellular proteins dependent on the cell context and surroundings (13, 21, 24). Therefore, we investigated the possible involvement of the PFP complex in mediating extracellular protein nutrient uptake under starvation for cell energy supply and survival.

Ovalbumin-DQ (OVA-DQ) is a fluorescent indicator that fluoresces upon proteolytic degradation (33). First, we used extracellular OVA-DQ to determine the possible βγ-CAT-driven extracellular protein uptake and intracellular degradation. After treating toad liver cells with 100 nM βγ-CAT, a striking increase in OVA-DQ fluorescence under Glc^-^/Gln^-^ conditions was observed by scanning confocal microscopy (Fig. 3A) and flow cytometry (Fig. 3B). Furthermore, the augmented fluorescence of OVA-DQ under Glc^-^/Gln^-^conditions was attenuated by immunodepletion of endogenous βγ-CAT (Fig. 3A and 3B). Consistent with the results in toad liver cells, the addition of βγ-CAT (40 nM) to starved mammalian HepG2 cells also promoted uptake and degradation of OVA-DQ (Fig. S3A). Furthermore, co-localization of βγ-CAT and OVA-DQ was readily observed, which exhibited a punctate pattern (Fig. 3C). Ethyl-isopropyl amiloride (EIPA) is a macropinocytosis inhibitor by inhibiting actin polymerization (34). The increase in extracellular protein (ovalbumin) internalization promoted by βγ-CAT was inhibited in both toad liver cells (Fig. S3B) and mammalian HepG2 cells (Fig. S3C) in the presence of the inhibitor. These results suggested that βγ-CAT mediated extracellular protein intake in starved cells by inducing macropinocytosis. Finally, βγ-CAT also enhanced internalization of toad *B. maxima* serum albumin (Bm-SA) under Glc^-^/Gln^-^ conditions (Fig. S3D). Collectively, these results demonstrated that βγ-CAT enhances uptake and intracellular degradation of extracellular proteins under cell starvation.

**Figure 3.**
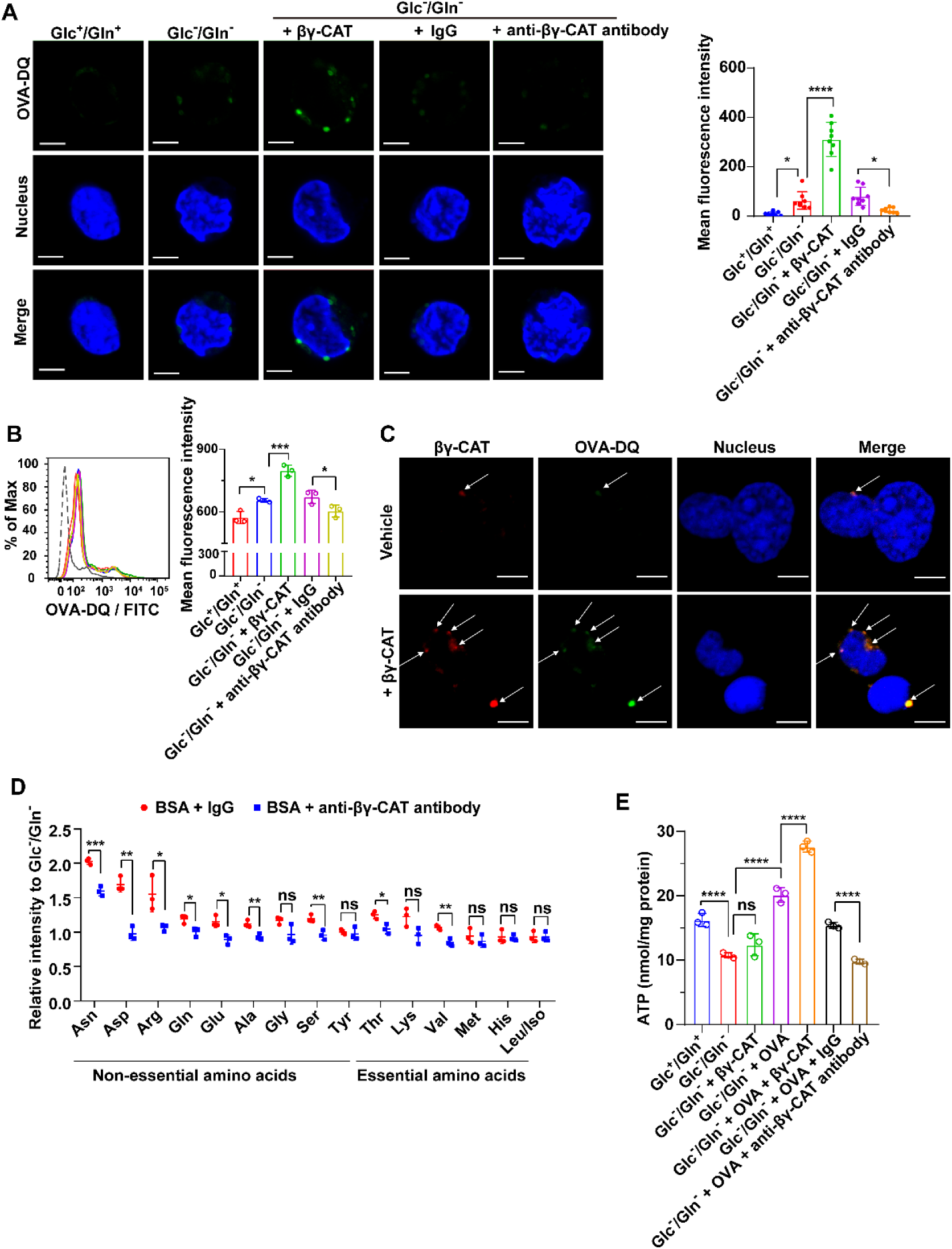
βγ-CAT promotes extracellular protein intake under cell starvation. (A, B) βγ-CAT augmented Ovalbumin-DQ intake in toad liver cells for energy supply under nutrient deficiency. Isolated toad liver cells were incubated with 100 nM βγ-CAT or 100 μg/mL anti-βγ-CAT antibodies in Glc^-^/Gln^-^ medium for 30 minutes. Then, the liver cells were incubated with 100 μg/mL (A) or 20 μg/mL (B) Ovalbumin-DQ (green) for 15 minutes, respectively. Ovalbumin-DQ fluorescence was determined by laser scanning confocal microscopy (A) and flow cytometry (B). Scale bars, 5 μm. The fluorescence intensity was quantified as described in (A) with eight images of each group (*Right*). (C) Colocalization (white arrow) of βγ-CAT and Ovalbumin-DQ in toad liver cells after incubation with 100 μg/mL Ovalbumin-DQ and 100 nM βγ-CAT in Glc^-^/Gln^-^ medium for 15 minutes. Scale bars, 5 μm. (D) Intracellular amino acid content of toad liver cells was determined by LC-MS and LC-MS/MS after incubation with 500 μg/mL BSA and 100 μg/mL anti-βγ-CAT antibodies in Glc^-^/Gln^-^ medium for 7 hours. (E) The ATP content in toad liver cells was determined by an ATP detection kit after incubation with 500 μg/mL OVA and 100 nM βγ-CAT or 100 μg/mL anti-βγ-CAT antibodies in Glc^-^/Gln^-^ medium for 7 hours. Rabbit IgG was used as an antibody control. Results (A, B, D, E) are reported as the mean ± SD of triplicate samples, ns (*P*≥0.05), \**P*<0.05, \*\**P*<0.01, \*\*\**P* <0.001, and \*\*\*\**P*<0.0001 by the one-way ANOVA (A, B, E) or unpaired *t-*test (D). All data are representative of at least two independent experiments.

Next, we further analyzed the metabolic situation of toad cells cultured in the presence of albumin under Glc^-^/Gln^-^ conditions for 7 hours by LC/MS and LC-MS/MS. The levels of amino acids, including asparagine and glutamine, in toad liver cells treated with bovine serum albumin (BSA) and βγ-CAT were augmented compared with those in cells treated with BSA only (Fig. S3E). Importantly, immunodepletion of endogenous βγ-CAT reduced the levels of several amino acids in toad liver cells, including threonine, valine, and nonessential amino acids such as asparagine, aspartic acid, arginine, glutamine, glutamic acid, alanine, and serine (Fig. 3D). These results revealed that βγ-CAT-mediated extracellular protein ingestion boosted the intracellular amino acid supply in toad cells under Glc^-^/Gln^-^ conditions.

To examine whether the increased amino acids were used for cell energy production, we analyzed total ATP availability in toad liver cells. Luciferase-based ATP assessment showed that ATP concentrations were significantly decreased under Glc^-^/Gln^-^ conditions, but partially recovered after addition of extracellular proteins (Figs. 3E and S3F). Under Glc^-^/Gln^-^ conditions, the addition of purified βγ-CAT to cultured toad cells augmented cellular ATP production in the presence of OVA (Fig. 3E) or BSA (Fig. S3F). Moreover, immunodepletion of endogenous βγ-CAT reduced ATP concentrations (Figs. 3E and S3F). Collectively, these results illustrated the capacity of βγ-CAT to drive extracellular protein import under cell starvation for cellular nutrient (amino acid) supply and ATP production.

### βγ-CAT supports toad cell survival in the presence of extracellular proteins under cell starvation

Because βγ-CAT was secreted by starved toad cells (Figs. 1 and 2), which mediates extracellular protein import and degradation, leading to increased amino acid supply and ATP production in starved toad cells (Fig. 3), it was reasonable to postulate that this PFP complex βγ-CAT could sustain the survival of starved cells in the presence of extracellular proteins.

The viability of toad cells was assessed by propidium iodide (PI) staining (35). In toad liver cells, no difference in cell viability among the various culture conditions at the zero-time point (stained immediately after the cells mixed with different media) was observed and approximately 90% of the cells were viable as analyzed by PI staining (Fig. S4A). However, toad liver cell death had increased substantially when the culture time was prolonged, and approximately 19% of liver cells had died after culture for 11 hours under Glc^+^/Gln^+^ conditions, while approximately 35% of liver cells had died under Glc^-^/Gln^-^ conditions (Fig. S4A). Notably, βγ-CAT alone did not obviously affect the survival rate of toad liver cells under starvation (Fig. S4A). However, the addition of βγ-CAT did increase the survival rate of starved toad cells in the presence of extracellular proteins BSA (Fig. 4A) or ovalbumin (OVA) (Fig. 4B). Additionally, immunodepletion of endogenous βγ-CAT greatly decreased the viability of starved toad cells (Fig. 4C and 4D). These results further verified the role of βγ-CAT in sustaining the survival of starved toad cells in the presence of extracellular proteins.

**Figure 4.**
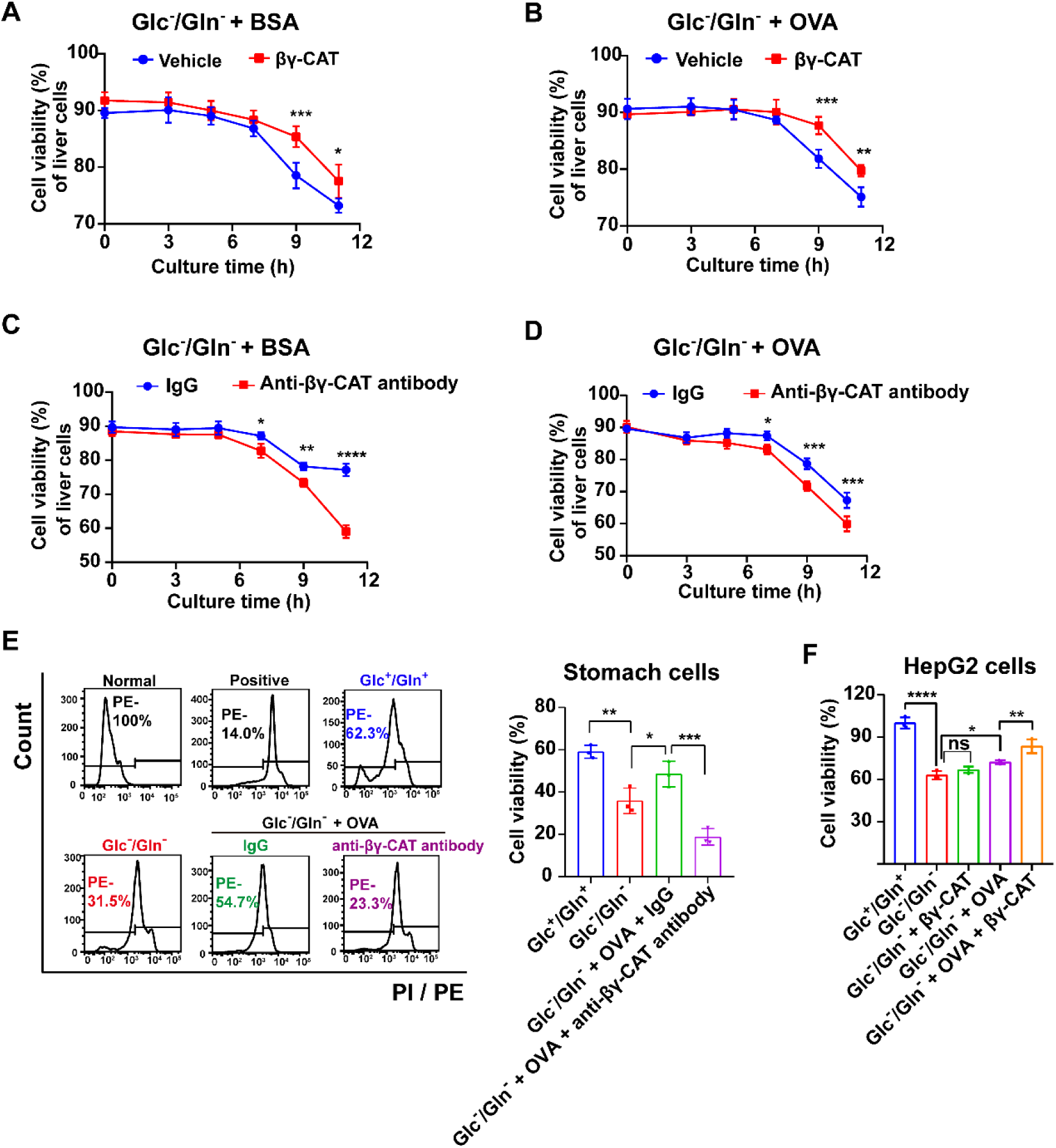
βγ-CAT supportes cell survival in the presence of extracellular proteins under cell starvation. (A, B) Addition of βγ-CAT promoted the viability of toad liver cells in the presence of extracellular proteins. Isolated toad liver cells were cultured with or without 100 nM βγ-CAT in the presence of 500 μg/mL BSA (A) or OVA (B) in Glc^-^/Gln^-^ medium for 0–11 hours. The cell viability was determined by PI staining. (C, D) Immunodepletion of endogenous βγ-CAT attenuated toad liver cell viability. Isolated toad liver cells were cultured with 100 μg/mL anti-βγ-CAT antibodies in the presence of 500 μg/mL BSA (C) or OVA (D) in Glc^-^/Gln^-^ medium for 0–11 hours. The cell viability was determined by PI staining. Rabbit IgG was used as an antibody control. (E) The viability of toad stomach cells was determined by PI staining after culture with 500 μg/mL OVA in the presence of 100 μg/mL anti-βγ-CAT antibodies in Glc^-^/Gln^-^ medium for 7 hours. Normal, untreated cells. Positive, cells treated with 0.5% Triton X-100 in Glc^-^/Gln^-^ medium for 5 minutes. IgG, rabbit antibody control. Right shows the quantitative comparison of cell viability in (E). (F) The viability of HepG2 cells was determined by MTS assays after culture in the presence of 5 mg/mL OVA with or without 40 nM βγ-CAT in Glc^-^/Gln^-^ medium for 36 hours. Results are reported as the mean ± SD of triplicate samples, ns (*P*≥0.05), \**P*<0.05, \*\**P*<0.01, *** *P*<0.001, and \*\*\*\**P*<0.0001 by two-way ANOVA (A–D) or one-way ANOVA (E, F). All data are representative of at least two independent experiments.

The ability of βγ-CAT to support the survival of starved toad cells in the presence of extracellular proteins was also determined in toad stomach epithelial cells. As expected, cell starvation increased PI-positive toad stomach cells after culture for 7 hours (Fig. 4E), an indication of a reduced cell survival rate. Consistent with the results of toad liver cells, immunodepletion of endogenous βγ-CAT with anti-βγ-CAT antibodies substantially reduced the survival rate of toad stomach cells in the presence of OVA (Fig. 4E). The protective effect of βγ-CAT was replicated in mammalian liver cell line HepG2. βγ-CAT (40 nM) showed no cytotoxicity in cells under Glc^-^/Gln^-^ conditions (Fig. S4B). The presence of βγ-CAT (40 nM) increased the survival rate of starved HepG2 cells in the presence of OVA (Fig. 4F).

Taken together, these results demonstrated the capacity of βγ-CAT to support cell survival in the presence of extracellular proteins under cell starvation.

### βγ-CAT is negatively regulated by high concentration of ATP *in vitro*

The above experimental evidence showed that toad cells secreted the PFP complex βγ-CAT under nutrient deficiency to scavenge extracellular proteins for their nutrient supply, ATP production, and survival (Figs.1, 3 and 4). Accordingly, such a cellular macromolecular nutrient acquisition and intracellular digestion system should be tightly regulated. In addition to the positive regulation of βγ-CAT expression and secretion by AMPK signaling (Fig. 2), there should be specific negative feedback regulators of the βγ-CAT-pathway. ATP, an end product of βγ-CAT actions (Fig. 3), might be a suitable candidate.

βγ-CAT oligomerizes and forms channels (pores) on liposomes, which induces dye release from lipid vesicles, an advantageous model without ATP receptors (12, 36). Interestingly, the dye release due to βγ-CAT channel formation on liposome was inhibited by 2.5 mM ATP or ADP, but not by AMP (Fig. 5A). The ATP concentration (2.5 mM) used was close to its physiological concentration in normal cells (5–10 mM) (37, 38). The inhibition of dye release through βγ-CAT channels by ATP (0.625–2.5 mM) was dependent on the concentration (Fig. 5B). Furthermore, a direct interaction between βγ-CAT and ATP was observed by a surface plasmon resonance (SPR) assay with an apparent *K*_*d*_ of approximately 2.8 × 10^−4^ M under our assay conditions (Fig. 5C). Notably, oligomer formation of βγ-CAT on liposome was not obviously changed in the presence of various ATP concentrations (0.625–2.5 mM) (Fig. 5D). This result suggested that ATP at these concentrations did not obviously affect oligomerization or membrane insertion of βγ-CAT. Finally, no ATPase activity was detected using purified βγ-CAT even with a dosage up to 1 μM (Fig. 5E). Collectively, these results revealed a direct interaction of ATP with βγ-CAT and suggested that high concentrations of ATP (1–5 mM) may negatively regulate the opening state of βγ-CAT channels.

**Figure 5.**
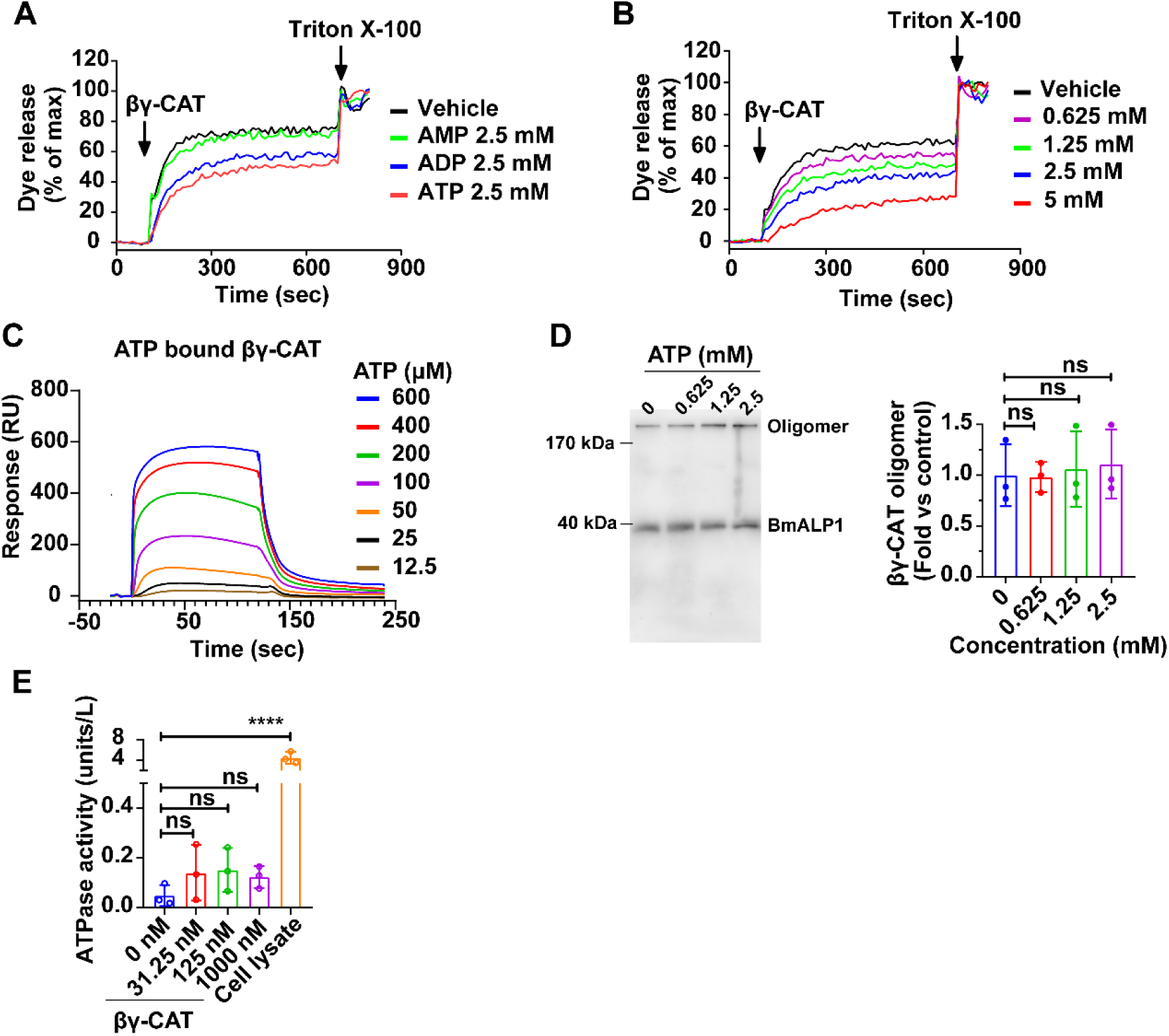
βγ-CAT is negatively regulated by high concentration of ATP *in vitro*. (A) Inhibitory effect of various adenosine phosphates (AMP, ADP, and ATP) on βγ-CAT-induced liposome dye release. (B) Concentration-dependent inhibitory effect of ATP (0–5 mM) on βγ-CAT-induced liposome dye release. (C) The direct interaction between βγ-CAT and ATP was analyzed by Biacore S200. (D) The effect of ATP on βγ-CAT oligomerization in liposomes was determined by western blotting (*Left*) and the bands of βγ-CAT oligomer were semi-quantified by ImageJ (*Right*). ATP was omitted as a control. (E) The potential ATPase activity of βγ-CAT was assayed by an ATPase activity kit. Data (D, E) are reported as the mean ± SD of triplicate samples, ns (*P*≥0.05) and \*\*\*\**P*<0.0001 by one-way ANOVA. All data are representative of at least two independent experiments.

## Discussion

PFPs are widely distributed in all kingdoms of life, which have long been recognized as either pore-forming toxin for microbial infection or host immune executors (11, 39, 40). Specially, the knowledge of these PFPs derived from animals and plants is mainly focused on their roles in cell death. The present study reported the direct and necessary function of PFP protein complex βγ-CAT in cell nutrient scavenging and energy supply under the deprivation of glucose and glutamine, two essential nutrient components in cell metabolism (2). We proposed an action model of βγ-CAT in driving the uptake of extracellular macromolecules such as proteins for cell nutrient supply and survival under starvation (Fig. 6). These findings provide further experimental evidence to support our previous hypothesis that βγ-CAT indeed acts as a novel cell vesicular delivery system which should play a physiological role in cell nutrient acquisition by mediating cellular nutrient import through endolysosomal pathways (13, 23, 24).

**Figure 6.**
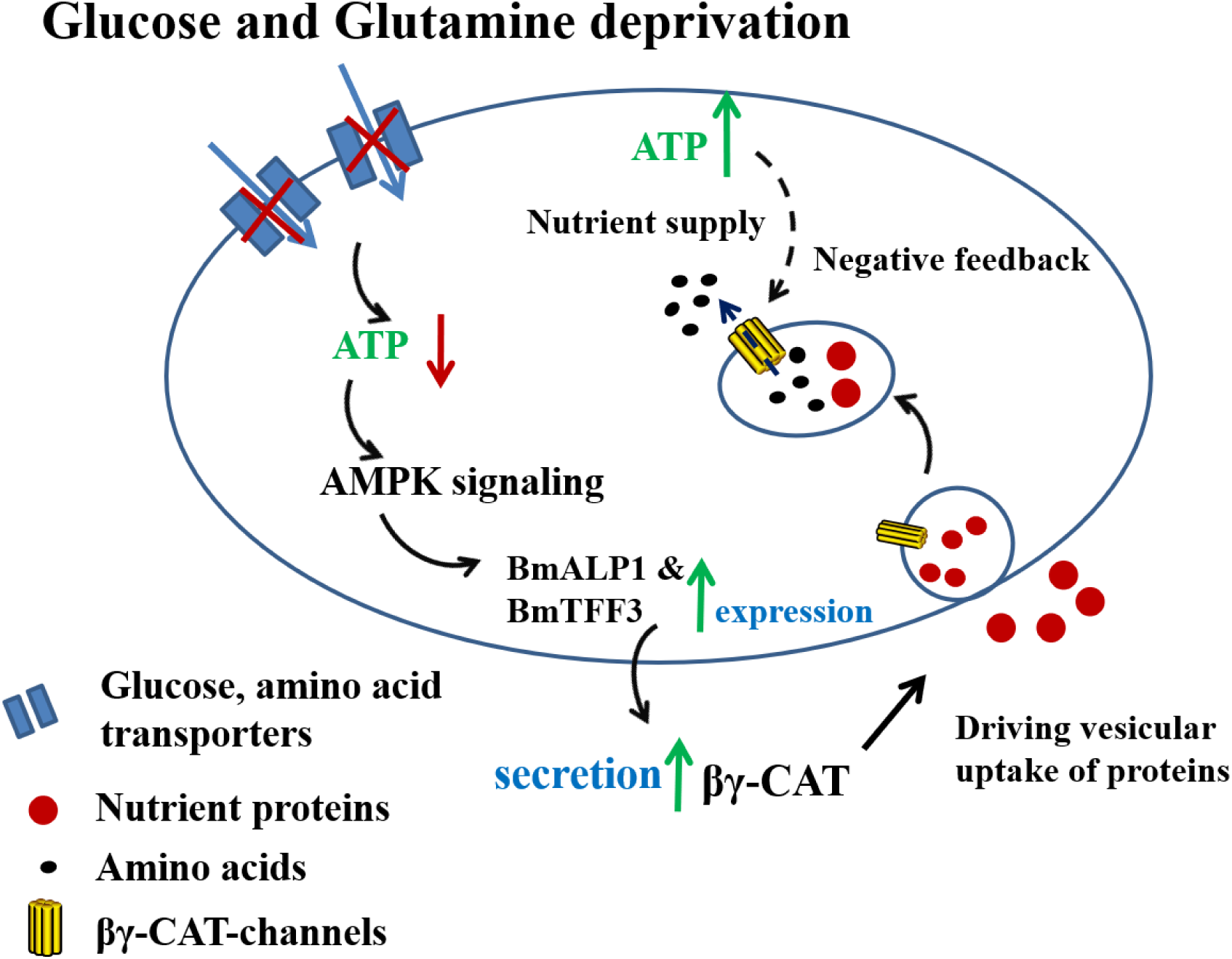
Proposed action model of PFP complex βγ-CAT in cell nutrient scavenging. Under Glc^-^/Gln^-^ conditions, AMPK signaling senses the deficiency of energy and nutrients and positively regulates the expression and secretion of βγ-CAT. βγ-CAT drives vesicular import of extracellular proteins through endolysosomal pathways probably by inducing pinocytosis/macropinocytosis. The acquired protein nutrients are hydrolyzed in endolysosomal organelles to facilitate intracellular amino acid supply and ATP production, thereby supporting cell survival. As a negative feedback, high concentrations of ATP (around 1–5 mM) may negatively regulate βγ-CAT channel opening (dashed black arrow).

Although constitutively expressed in toad cells in nutrient-rich medium (Fig. 1), βγ-CAT was not secreted into the culture supernatant, indicating that the cells could acquire nutrients through membrane transporters under normal nutrient-rich environments. Under Glc^-^/Gln^-^ conditions, the attenuated synthesis of βγ-CAT in toad cells at the beginning of starvation (Fig. 1A and 1B) was in accordance with a fact that cells decrease or even stop their protein production during nutrient deficiency (41, 42). However, the expression of βγ-CAT was increased again afterwards. Moreover, βγ-CAT secreted into the culture supernatant was continuously augmented, which was readily attenuated by addition of glucose to the medium (Fig. 1). Therefore, these results clearly illustrate that βγ-CAT is a protein machine dispatched by toad cells in response to nutrient deficiency.

Intracellular nutrient sensors, which survey the abundance of energy and major metabolites, play an important role in metabolic homeostasis and cell survival (1, 2). AMPK signaling is involved in sensing nutrient and energy availability, which is switched on by a lack of energy or nutrients (43, 44). Our assays employing pharmacological inhibitors suggested that AMPK signaling controls the expression and secretion of βγ-CAT under cell nutrient deficiency (Fig 2). These results are in accordance with the role of AMPK signaling in regulating cell responses to nutrient and energy deficiency. The present observations revealed that starved toad cells secrete a PFP machine such as βγ-CAT downstream of AMPK signaling to scavenge and mediate cellular uptake of extracellular proteins for cell energy supply and survival. This PFP-driven vesicular delivery of extracellular nutrients represents an unknown strategy and mechanism involved in cell nutrient acquisition. Our data suggest that AMPK signaling is a positive regulator of the βγ-CAT pathway, but the detailed molecular mechanisms are unclear at present stage, which is an intriguing question in future study.

Diverse endocytic pathways are available at the surface of metazoan cells (3). Macropinocytosis is an actin-dependent endocytic pathway that produces endocytic vesicles (>200 nm) (45, 46), whereas pinocytosis refers to cell internalization of extracellular fluid into small endocytic vesicles (<200 nm) (3, 7). In a murine DC model, βγ-CAT enhances pinocytosis as determined by uptake of Lucifer Yellow, which enhances import of the antigen OVA (21). βγ-CAT also drives macropinocytosis *in vivo* and *in vitro* in toad osmoregulatory organs to facilitate toad water maintaining (13). Consistently, pharmacological inhibitors in the present study also suggested that βγ-CAT promoted uptake of extracellular proteins via pinocytosis/macropinocytosis (Figs. 3 and S3). However, it is worth pointing out that the endocytic form mediating protein uptake in toad cells, including liver and gastrointestinal cells, stimulated by βγ-CAT has not been completely clarified, especially *in vivo*. It is possible that βγ-CAT may employ multiple cell entry mechanisms to scavenge protein nutrients depending on the cell context and surroundings, which is worthy of further investigation.

The functional diameter of bacterial toxin aerolysin channels is approximately 1.5 nm, which is large enough for translocation of oligonucleotides, peptides, and unfolded proteins (47, 48). As a member of af-PFPs, βγ-CAT forms channels on membranes with a functional diameter similar to that of aerolysin (12-14). Previously, it has been observed that the channels formed by βγ-CAT served for the translocation of processed OVA peptides to cytosol for antigen presentation in murine DC cells (21). Accordingly, the βγ-CAT channels formed on endolysosomes of toad cells may act as channels to traverse amino acids and small peptides produced from imported proteins by proteolytic hydrolysis to cytosol for cell nutrient supply and energy (Fig. 3).

The physiological concentrations of ATP in cells are estimated to be 5–10 mM (37, 49). Extracellular ATP levels are at the micromolar level (50–200 µM) under pathological conditions such as a tumor microenvironment, whereas in healthy tissues, extracellular ATP concentrations are sub-micromolar (likely about 10–100 nM) (38). βγ-CAT did not possess an ATP-hydrolyzing activity (Fig. 5E). In liposome assays, the presence of ATP did not obviously affect oligomerization of βγ-CAT, but dye release from βγ-CAT channels formed in liposome was inhibited in a concentration-dependent manner by ATP (0.6–5 mM), but not by AMP. These results suggested that the normal intracellular contents of ATP could negatively regulate the opening state of βγ-CAT channels (Fig. 5). In another word, βγ-CAT may sense ATP abundance to regulate the opening state of its channels. ATP is an energy situation marker in cells. The negative regulation of βγ-CAT channels by high concentrations of ATP (1–5 mM) is inconsistent with the fact that this PFP complex is secreted from starved cells to promote uptake of extracellular nutrients in response to a poor energy status (Figs. 1 and 3). Furthermore, the concentration-dependent regulation of βγ-CAT channels by ATP may lead to transcellular delivery of nutrients to internal tissue environments such as tissue parenchymal cells by the release of nutrient exosomes (24).

βγ-CAT is a complex of BmALP1 and BmTFF3, in which BmTFF3 acts as a chaperon and regulatory unit of BmALP1 to stabilize the PFP monomer and deliver it to proper targets (16, 17). βγ-CAT undergoes oligomerization and forms channels on membranes. The inhibition of βγ-CAT channels by ATP indicated that ATP binds to the PFP complex, which is in accordance with the binding of ATP to βγ-CAT (Fig. 5C). However, at present, the exact binding sites of ATP on βγ-CAT and the regulatory mechanisms are unknown and important future research directions. The findings that βγ-CAT is positively regulated by AMPK signaling but negatively regulated by high concentration of ATP (1–5 mM) further emphasize and support the notion that the PFP complex is necessary for cell nutrient acquisition and metabolic flexibility.

It is well documented that autophagy is a cellular process to sequester and degrade intracellular components, which play roles in cell responses to nutrient starvation (50, 51). Comparatively, the SELC protein βγ-CAT represents a novel cellular strategy to sense and uptake extracellular macromolecules like proteins as nutrients under cell starvation. This novel cell nutrient acquisition pathway mediated by a secretory PFP such as βγ-CAT through endolysosomal systems should be especially significant in the absence of essential small nutrient compounds, including glucose and glutamine, as revealed in the present study (Fig. 6). It may also be necessary and essential when classic plasma membrane-integrated transporters including solute carriers (SLCs) are absent such as in undifferentiated cells or they do not work properly. Cross-regulation of autophagy and macropinocytosis is poorly understood (8). The coordinative regulation between autophagy to sequeste intracellular components and PFP-driven cell uptake to import extracellular compounds like that mediated by βγ-CAT is worthy of further study.

SELC protein βγ-CAT works in cell nutrient acquisition at least at two levels. First, toad *B. maxima* cells secrete βγ-CAT to scavenge extracellular nutrients under nutrient deficiency at the cellular level (present study). Second, βγ-CAT in toad *B. maxima* blood circulation is an immediate and active responsive element under toad fasting *in vivo*, which transcellularly deliver and transport albumin-bound fatty acids to tissue parenchymal cells for their nutrient supply (24). These findings uncovered the primary and necessary role of this PFP machine in toad *B. maxima* physiology for adaptation to various nutrient environments. Rationally, similar strategies and executive pathways should be conserved in vertebrates, in which various families of PFPs including af-PFPs are widely distributed. Knowledge from βγ-CAT can provide clues to understand novel PFP-driven cell vesicular delivery systems in nutrient acquisition and metabolic flexibility. Although af-PFPs have not been clearly observed in Eutherian mammals, other PFP family members can readily compensate for the role of af-PFPs.

In conclusion, the present study elucidated that toad *B. maxima* cells secrete βγ-CAT, a PFP and TFF complex assembled depending on environmental cues under glucose and glutamine deficiency. This PFP complex supports cell survival by driving the cellular import of extracellular proteins through endolysosomal pathways. The imported proteins serve as nutrients in starved cells for energy supply. AMPK signaling positively regulates the expression and secretion of βγ-CAT, whereas high concentrations of ATP (>1 mM) bind to and negatively regulate βγ-CAT channels. Our findings define the essential role of a PFP in cell macromolecular nutrient scavenging, providing a new paradigm for PFPs in cell nutrient acquisition and metabolic flexibility.

## Acknowledgements

We thank Dr. Lian Yang (Kunming Institute of Botany, Chinese Academy of Sciences) for her help with the interaction detection by using SPR S200. We are grateful to Dr. Lin Zeng (Kunming Biological Diversity Regional Center of Instrument) for their assistances in LC/MS and LC/MS/MS analysis. This work was supported by grants from the National Natural Science Foundation of China (grant numbers 31572268, U1602225, and 31872226) and the Yunling Scholar Program to Yun Zhang.

## Author contributions

Y.Z., L.Z.L, L.L. and Y.X conceived of and conceptualized the study; L.Z.L., L.L. Z.H.S, X.L.B and Q.Q.W did the experiments, L.Z.L, Y.Z., L.L. and Y.X analyzed and interpreted the data; L.Z.L, L.L., Y.Z. and Y.X. wrote the manuscript; Y.Z., L.Z.L., L.L., Z.H.S, X.L.B and Q.Q.W critically revised the manuscript for important intellectual content.

## Materials and Methods

### Animal

Feeding of toads (*B. maxima*) was performed as described previously (19). All procedures and the care and handing of animals were approved by the Ethics Committee of the Kunming Institute of Zoology, Chinese Academy of Sciences (Approval ID: IACUC-OE-2021-05-001).

### Cell culture

Toad liver cells were isolated by a two-step EDTA/collagenase perfusion technique as previously described with the following modifications (52). The perfusion solution Ringer’s buffer and perfusion solution II containing 1 mg/mL collagenase (Solarbio, Cat C8140) were used for toad liver tissue perfusion. After perfusion, liver tissues from three toads were cut into pieces and washed with Ringer’s buffer once and then oscillatory digested in perfusion solution II at 26°C for 1 hour. The cells were filtered through a 40-μm mesh and collected by centrifugation at 805 g for 5 minutes at 4°C. Toad stomach and intestinal cells were isolated as described previously (13).

The mammalian cell line HepG2 was purchased from Kunming Cell Bank, Chinese Academy of Sciences. Cells were cultured in DMEM/F-12 (Biological Industries, Cat 01-172-1A) containing 10% fetal bovine serum (Biological Industries, Cat 04-001-1A) and 1% Penicillin-Streptomycin Solution (Biological Industries, Cat 03-031-1B) at 37°C with 5% CO_2_.

### Glucose and glutamine deprivation and treatment

Cells were cultured in basal medium containing 3.151 g/L glucose and 2.5 mM glutamine (Glc^+^/Gln^+^ medium, Biological Industries, Cat 01-172-1A), glucose and glutamine-free medium (Glc^-^/Gln^-^ medium, Biological Industries, Cat 01-057-1A), or 3.151 g/L glucose but glutamine-depleted medium (Glc^+^/Gln^-^ medium). For macropinocytosis inhibition, cells were first incubated with 100 μM EIPA (MedChemExpress, Cat HY-101840) in Glc^-^/Gln^-^ medium for 1 hour at 26°C (toad cells) or 37°C (HepG2 cells), respectively. For AMPK signaling inhibition, cells were treated with 0–10 μM compound C or 0–20 μM SBI-0206965 (MedChemExpress, Cat HY-13418 and Cat HY-16966, respectively) in Glc^-^/Gln^-^ medium for 3 hours at 26°C. To deplete endogenous βγ-CAT, toad cells were incubated with 100 μg/mL anti-βγ-CAT rabbit polyclonal antibodies or 100 μg/mL rabbit IgG (Proteintech, Cat B900610) as the isotype control in Glc^-^/Gln^-^ medium at 26°C.

### Cell viability assays

Cell viability of toad cells was measured using propidium iodide (PI) stain assay as described previously (35). A total of 5×10^6^ isolated toad liver cells were cultured at 26°C in various media for 0, 3, 5 and then up to 11 hours at 2-hour intervals. The treatments were as follows: Glc^+^/Gln^+^ medium, Glc^-^/Gln^-^ medium, Glc^-^/Gln^-^ medium containing 100 nM βγ-CAT, Glc^-^/Gln^-^ medium containing 500 μg/mL ovalbumin (OVA, Coolaber, Cat CA1421) or BSA (Sigma-Aldrich, Cat A9418), Glc^-^/Gln^-^ medium containing 100 nM βγ-CAT and 500 μg/mL OVA or BSA, Glc^-^/Gln^-^ medium containing 100 μg/mL anti-βγ-CAT antibodies in the presence of 500 μg/mL OVA or BSA, and Glc^-^/Gln^-^ medium containing 100 μg/mL rabbit IgG in the presence of 500 μg/mL OVA or BSA. To assess the viability of toad stomach cells, 5×10^6^ isolated toad stomach cells were subjected to various treatments (Glc^+^/Gln^+^ medium, Glc^-^/Gln^-^ medium, and Glc^-^/Gln^-^ medium containing 500 μg/mL OVA with rabbit IgG or anti-βγ-CAT antibodies) for 7 hours at 26°C. To assess cytotoxicity of compound C and SBI-0206965, 5×10^6^ isolated toad liver cells were cultured in Glc^-^/Gln^-^ medium plus compound C (0–10 μM) or SBI-0206965 (0–20 μM) for 3 hours at 26°C. After treatments, toad cells were stained with 500 ng/mL PI (ACMEC, Cat P72320) for 4 minutes at 26°C. Fluorescence was recorded using a LSR Fortessa cell analyzer (Becton Dickinson, Franklin Lakes, NJ, USA).

To assess viability of mammalian HepG2 cells, 1×10^4^ cells were seeded into each well of a 96-well culture plate. After overnight culture, the cells were washed thrice with PBS and incubated in Glc^+^/Gln^+^ medium, Glc^-^/Gln^-^ medium, Glc^-^/Gln^-^ medium plus various concentrations of βγ-CAT (0–160 nM), or Glc^-^/Gln^-^ medium plus 5 mg/mL OVA in the presence or absence of 40 nM βγ-CAT for 36 hours. Then, the cells were incubated with MTS reagent (Promega, Cat G3580) for 2 hours in the dark at 37°C with 5% CO_2_. Absorbance was read at 490 nm with an Infinite 200 Pro microplate reader (Tecan, Männedorf, Switzerland).

### Hemolytic activity assay

A total of 1×10^7^ toad liver, intestinal, or stomach cells was collected and then washed twice with 30 mL Ringer’s solution. Toad liver cells were cultured in Glc^+^/Gln^+^, Glc^-^/Gln^-^, or Glc^+^/Gln^-^ medium for 1–5 hours at 26°C, and intestinal or stomach cells was cultured for 1 hour at 26°C. Then, culture supernatants were collected and prepared for the hemolytic activity assay as described previously (13). Maximum hemolysis (100% lysis) was defined as 0.5% Triton X-100-lysed samples in culture medium. To determine whether hemolytic activity was induced by βγ-CAT, the culture supernatant of toad liver cells was mixed with 100 μg/mL anti-βγ-CAT antibodies or 100 μg/mL rabbit IgG as the control, and then hemolytic activity was tested.

### Extracellular protein intake and degradation

A total of 5×10^6^ isolated toad liver cells were treated as described in the “**Glucose and glutamine deprivation and treatment**” section at 26°C for 30 minutes. Then, the cells incubated with 100 μg/mL FITC-OVA (Bioss, Cat bs-0283P-FITC), FITC-Bm-SA (FITC-labeled toad *B. maxima* serum albumin (24)) or 20 μg/mL Ovalbumin-DQ (ThermoFisher, Cat D12053) in the dark at 26°C for 15 minutes. Fluorescence was detected by the LSR Fortessa cell analyzer using the FITC channel. In each sample, 1×10^4^ single cells were analyzed.

For immunofluorescence, cells were incubated with 100 μg/mL Ovalbumin-DQ in the dark at 26°C for 15 minutes. To stain βγ-CAT, cells were washed and incubated with 20 μg/mL anti-βγ-CAT primary antibodies overnight at 4°C in the dark. After washing with PBS three times, the cells were incubated with 10 μg/mL cy3-conjugated anti-rabbit IgG for 1 hour at 37°C in the dark. Then, the samples were sealed with an anti-fluorescent quench agent containing DAPI. Images were acquired by a Zeiss LSM 880 microscope system (Carl Zeiss, Oberkochen, Germany).

### Amino acid analysis

A total of 1×10^7^ isolated toad liver cells were treated in various media as described in the “**Cell viability assays**” section for 7 hours at 26°C. Amino acid analysis was performed as previously described using mass spectrometry-based methods (33). After harvesting, cells were washed twice with cold 150 mM ammonium acetate solution (pH 7.4, 4°C). Amino acids were extracted with a ddH_2_O and MeOH solution (1:4) at -80°C, and then the samples were incubated for 15 minutes at -80°C. Then, the supernatants were collected by centrifuging at 17,000 g for 10 minutes. 10 μL of these supernatants were injected per analysis. Samples were run on a Vanquish (Thermo Fisher Scientific) UHPLC system with mobile phase A (5 mM NH_4_AcO, pH 5.0) and mobile phase B (ACN) at a flow rate of 200 μL/min on an Accucore™-150-Amide-HILIC column 2.6 μm (100 × 2.1 mm) (Thermo Fisher Scientific, Cat 16726-102130) at 40 °C at a gradient from 45% to 90% A in 15 minutes followed by a 10-minute isocratic step. The UHPLC was coupled to a Q-Exactive (Thermo Fisher Scientific) mass analyzer running in polarity switching mode at 3.5 kV for positive change scanning and 3.0 kV for negative change scanning at an MS1 resolution of 70,000. Metabolites were identified by exact mass (MS1), retention time, and in some cases, by their fragmentation patterns (MS2) at normalized collision energy. Quantification was performed by area under the curve integration of MS1 ion chromatograms with the Thermo Scientific Xcalibur software package. Area values were normalized to cell count averages from triplicate wells treated in parallel for each condition.

### ATP measurement

Intracellular ATP was extracted and measured by an ATP detection kit (Beyotime, Cat S0026) in accordance with the manufacturer’s protocol. Briefly, toad liver cells were treated same as described in the “**Cell viability assay**” section and harvested at 7 hours. After rinsing in cold glucose-free Ringer’s buffer twice, cell pellets were lysed in ice-cold lysis buffer and centrifuged at 10,625 g for 5 minutes at 4°C, and then the supernatants were subjected to Luminoskan Ascent detection (Tecan, Männedorf, Switzerland).

### Liposome dye release assay

To assess the effect of adenosine phosphate on βγ-CAT channel formation-induced liposome dye release, liposomes were incubated with 600 nM βγ-CAT in the presence or absence of adenosine phosphate (AMP, ADP, and ATP) (Sigma), and then dye release assays were performed as described previously (12).

### Purification of βγ-CAT

The purification of βγ-CAT were performed as described previously (14).

### Surface plasmon resonance interaction analysis (BIAcore)

A direct interaction was assessed as described previously (53). Briefly, purified βγ-CAT was diluted in 10 mM sodium acetate (pH 5.0) at 20 μg/mL, and 20000 response units were immobilized via amine coupling to CM7 sensor chip flow chambers (Cytiva, Cat 28953828) in accordance with the manufacturer’s instructions in a BIAcore S200 instrument. A flow chamber subjected to the immobilization protocol, but without any addition of protein, was used as a control (blank). All interaction experiments were conducted at 25°C in normal saline. βγ-CAT associated with ATP for 120 seconds, and the dissociation time was 120 seconds. Binding curves were displayed, and the dissociation (K_*d*_) rate constants were determined using BIA evaluation 4.1 software and its equation for 1:1 Langmuir binding.

### Western blotting

To assess protein level of βγ-CAT in toad liver cell lysate and supernatant, cells were cultured in Glc^+^/Gln^+^, Glc^-^/Gln^-^, or Glc^+^/Gln^-^ medium, respectively. Then, the cells were lysed and prepared for western blotting. The cell supernatant was concentrated to one tenth of the original volume by a vacuum lyophilizer for western blotting as described previously (16).

To assess AMP-activated protein kinase (AMPK) signaling, cells were treated as described in the “**Glucose and glutamine deprivation and treatment**” section. Then, the cells were lysed and subjected to immunoblotting as described above. An anti-βγ-CAT polyclonal antibody (14), anti-AMPKα1/2 (Bioworld, Cat BS1009), anti-AMPKα1/2 (phospho-T183/172) (Bioworld, Cat BS5003), anti-ACC1 (Affinity Biosciences, Cat AF6421), anti-ACC1 (phospho-S79) (Affinity Biosciences, Cat AF3421), and anti-β-actin (Proteintech, Cat 66009-1-Ig) primary antibodies were used in these experiments.

### Quantitative real-time PCR

The mRNA levels of BmALP1 (βγ-CAT-α) and BmTFF3 (βγ-CAT-β) in toad liver cells were measured by quantitative real-time PCR (qRT-PCR) using a ChamQ Universal SYBR qPCR Master Mix kit (Vazyme, Cat Q711-02). The cycle counts of target genes were normalized to those of β-actin. Primer sequences of βγ-CAT and β-actin used were the same as report previously (18).

### ATPase activity assay

The ATPase activity of βγ-CAT was measured by an ATPase activity assay kit (Sigma-Aldrich, Cat MAK113-1KT) according to the manufacturer’s protocol. Briefly, various concentrations βγ-CAT was incubated with 4 mM ATP in assay buffer (40 mM Tris, 80 mM NaCl, 8 mM MgAc_2_, and 1 mM EDTA, pH 7.5) for 30 minutes at 26°C. Cell lysates from HepG2 cells were used as a positive control. Absorbance was read at 620 nm, and the free phosphate concentration was calculated by a standard curve (0–50 μM free phosphate).

### Sequence alignment

Sequence alignment analysis of AMPKα and ACC1 in *B. maxima* with other species was conducted by Clustal Omega6 (54). The figure was generated using Jalview.

## Statistical analysis

All experimental values are expressed as means ± SD. Each experiment was repeated at least twice. All data were analyzed using GraphPad Prism 8.0 software. Two comparisons were performed using the standard unpaired *t*-test. Multiple comparisons were performed by one-way ANOVA with post-hoc contrasts by Dunnett’s multiple comparisons test. Multi-group comparisons were made by two-way ANOVA with Sidak’s multiple comparisons test. *P<*0.05 was considered statistically significant.

---

**Fig. S1.**
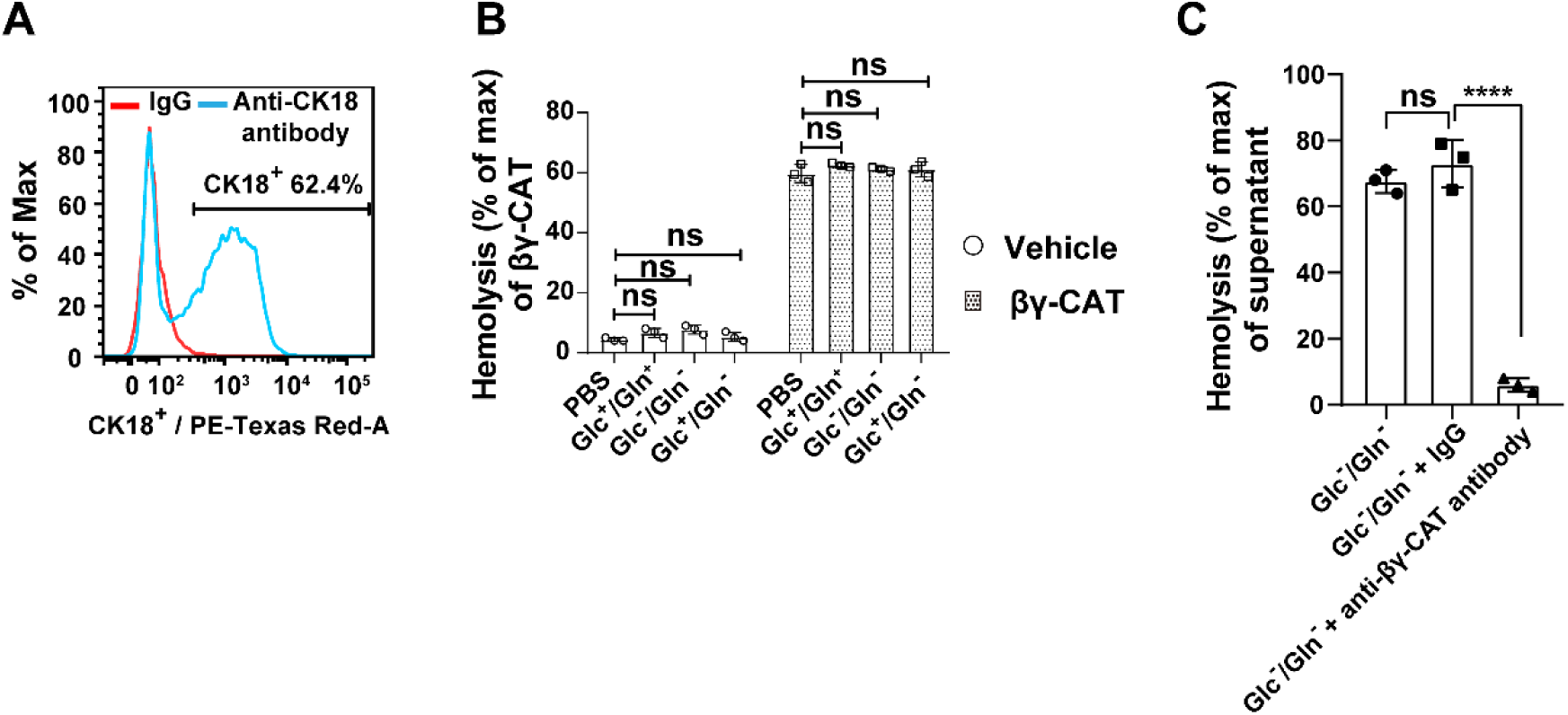
(A) The proportion of CK18^+^ hepatocytes was determined by flow cytometry with an anti-cytokeratin 18 (CK18, a hepatocyte marker) antibody. (B) The hemolytic activity of exogenous βγ-CAT (5 nM) was assayed in PBS, Glc^+^/Gln^+^, Glc^-^/Gln^-^, or Glc^+^/Gln^-^ medium, respectively. (C) The hemolytic activity of liver cell supernatants under Glc^-^/Gln^-^ conditions was detected in the presence of anti-βγ-CAT antibodies (100 μg/mL) or rabbit IgG (an antibody control). Data (B, C) are represented as the mean ± SD of triplicate samples. ns (*P*≥0.05) and \*\*\*\**P*<0.0001 by two-way ANOVA (B) or by one-way ANOVA (C). All data are representative of at least two independent experiments.

**Fig. S2.**
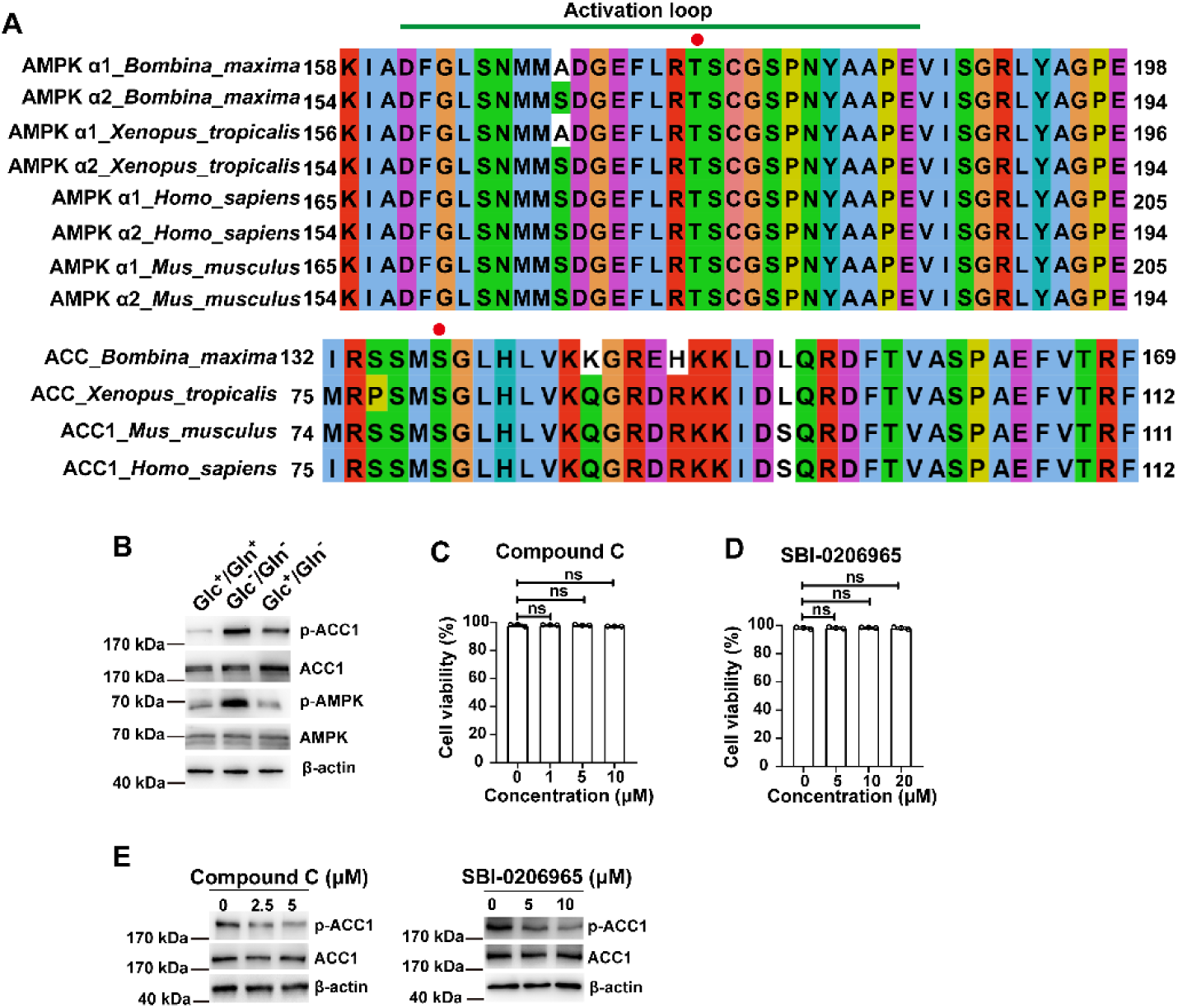
Compound C and SBI-0206965 inhibit activation of AMPK signaling in toad liver cells under Glc^-^/Gln^-^ conditions. (A) Sequence alignment of AMPKs (*Top*) and ACCs (*Bottom*) in toad *B. maxima* and other species was analyzed by Clustal Omega6. The activation loop of AMPK is indicated by a green line. The phosphorylated site (Thr residue of AMPKs and Ser residue of ACCs) is indicated by a red spot. (B) AMPK signaling activation in toad liver cells after culture in Glc^+^/Gln^+^, Glc^-^/Gln^-^, or Glc^+^/Gln^-^ medium for 3 hours was detected by western blotting. (C, D) Cytotoxicity of compound C (C) and SBI-0206965 (D) in toad liver cells was detected by PI staining after treatment with compound C or SBI-0206965 in Glc^-^/Gln^-^ medium for 3 hours. (E) Phosphorylation of pS79-ACC1 in toad liver cells was determined by western blotting after treatment with compound C or SBI-0206965 in Glc^-^/Gln^-^ medium for 3 hours. Data (C, D) are represented as the mean ± SD of triplicate samples. ns (*P*≥0.05) by one-way ANOVA. All data are representative of at least two independent experiments.

**Fig. S3.**
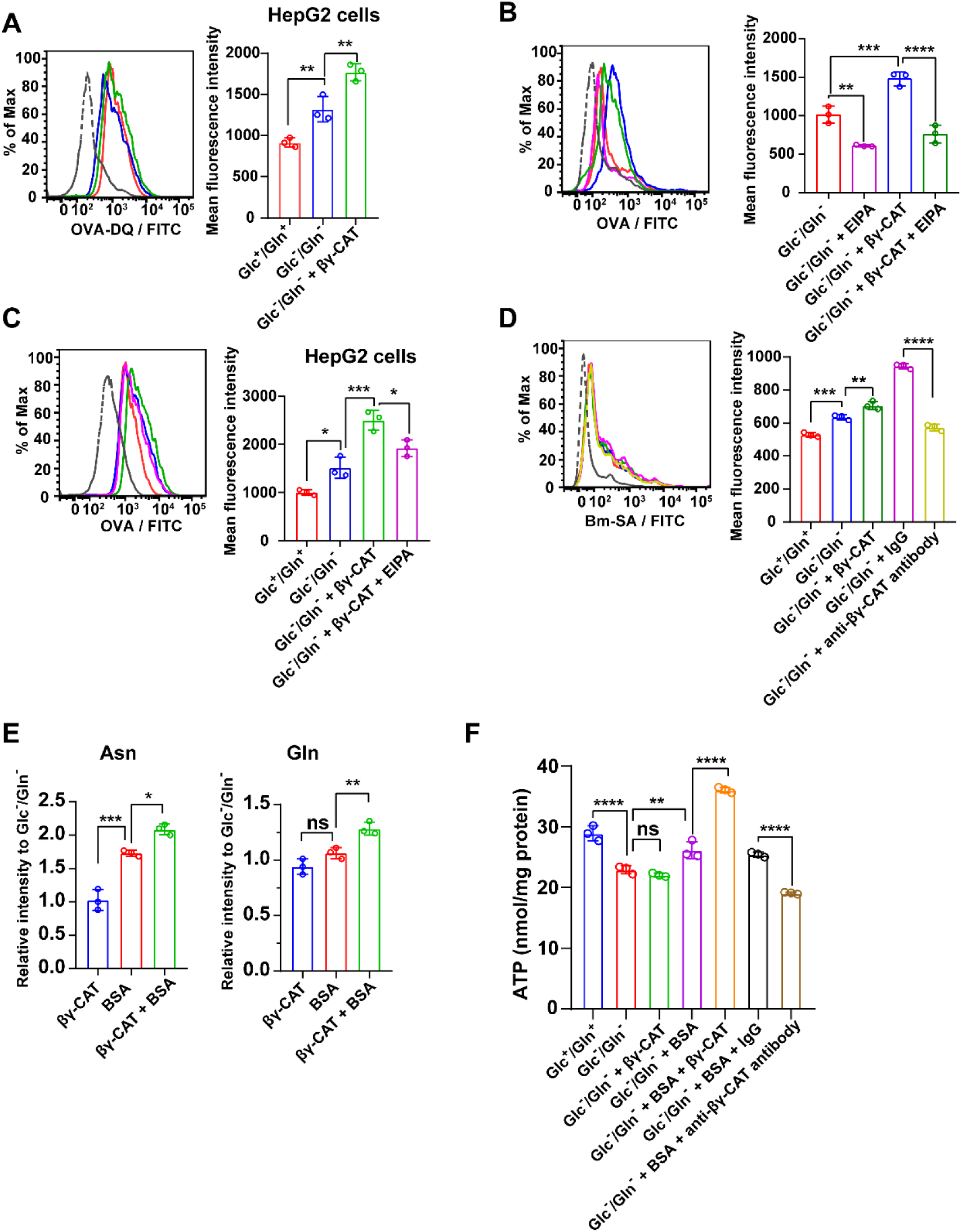
βγ-CAT promotes extracellular protein intake under cell starvation. (A) The mean fluorescence intensity of Ovalbumin-DQ in HepG2 cells was determined by flow cytometry after treatment with 40 nM βγ-CAT and 20 μg/mL Ovalbumin-DQ in Glc^-^/Gln^-^ medium for 30 minutes. (B, C) Inhibitory effect of a macropinocytosis inhibitor (EIPA) on protein intake induced by βγ-CAT. Toad liver cells (B) and HepG2 cells (C) were incubated with or without 100 μM EIPA in Glc^-^/Gln^-^ medium for 1 hour. Then, the cells were incubated with 100 μg/mL FITC-OVA in the presence of 100 nM βγ-CAT (toad liver cells) or 40 nM βγ-CAT (HepG2 cells) for 30 minutes. The mean fluorescence intensity of Ovalbumin-DQ in toad liver cells and HepG2 cells was determined by flow cytometry. (D) The mean fluorescence intensity of Bm-SA in toad liver cells was determined by flow cytometry after treatment with 100 nM βγ-CAT or 100 μg/mL anti-βγ-CAT antibodies in Glc^-^/Gln^-^ medium for 30 minutes. (E) The intracellular content of representative amino acids in toad liver cells was determined by LC-MS and LC-MS/MS after incubation with 500 μg/mL BSA and 100 nM βγ-CAT in Glc^-^/Gln^-^ medium for 7 hours. (F) The ATP content in toad liver cells was determined by an ATP detection kit after incubation with 500 μg/mL BSA and 100 nM βγ-CAT or 100 μg/mL anti-βγ-CAT antibodies in Glc^-^/Gln^-^ medium for 7 hours. Rabbit IgG was used as an antibody control. Results are reported as the mean ± SD of triplicate samples, ns (*P*≥0.05), \**P* <0.05, \*\**P*<0.01, \*\*\**P*<0.001, and \*\*\*\**P* <0.0001 by the one-way ANOVA. All data are representative of at least two independent experiments.

**Fig. S4.**
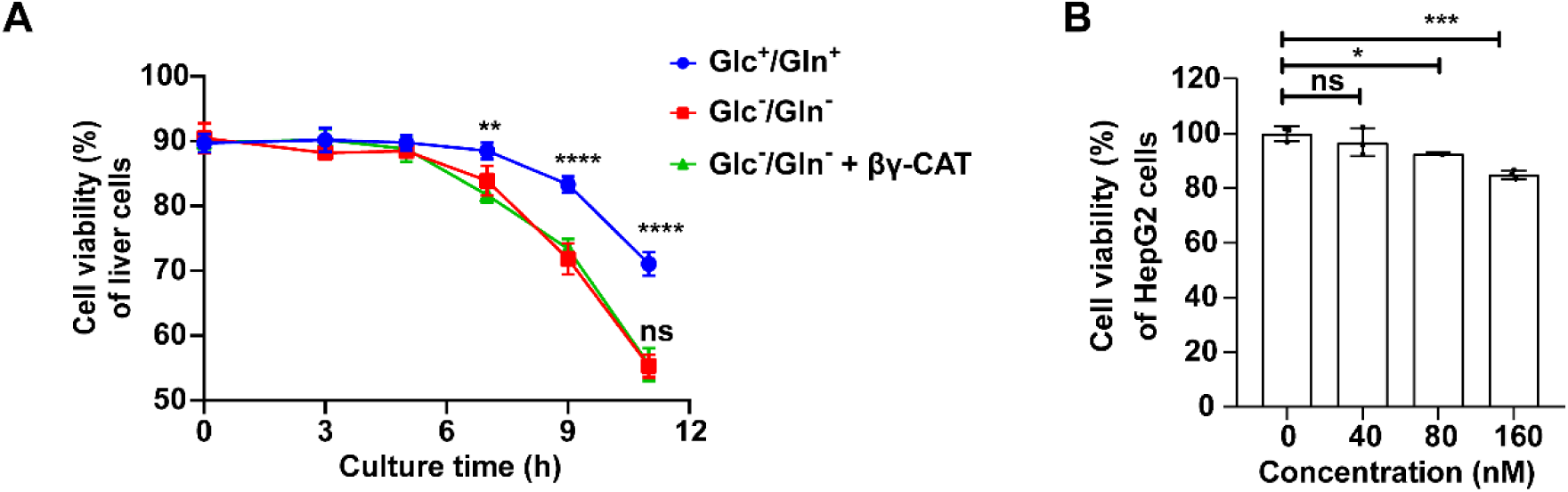
Viability of toad liver cells and cytotoxicity of βγ-CAT in HepG2 cells under Glc^-^/Gln^-^ conditions. (A) The viability of toad liver cells was determined by PI staining after culture in Glc^+^/Gln^+^, Glc^-^/Gln^-^, or Glc^-^/Gln^-^ plus βγ-CAT (100 nM) for 0–11 hours. (B) Cytotoxicity of βγ-CAT in HepG2 cells was detected by MTS assays after treatment with various concentrations of βγ-CAT in Glc^-^/Gln^-^ medium for 36 hours. Results are reported as the mean ± SD of triplicate samples. In (A), statistical significance was calculated by two-way ANOVA, \*\**P*<0.01, and \*\*\*\**P*<0.0001 (Glc^-^/Gln^-^ vs. Glc^+^/Gln^+^ groups) and ns *P*≥0.05 (Glc^-^/Gln^-^ + βγ-CAT vs. Glc^-^/Gln^-^ groups). In (B) ns (*P*≥0.05), \**P*<0.05, and \*\*\**P*<0.001 by one-way ANOVA. All data are representative of at least two independent experiments.

